# Stroma transcriptomic and proteomic profile of prostate cancer metastasis xenograft models reveals conservation of bone microenvironment signatures

**DOI:** 10.1101/2020.04.03.018143

**Authors:** S. Karkampouna, M.R. De Filippo, C.Y. Ng, I. Klima, E. Zoni, M. Spahn, F. Stein, P. Haberkant, G.N. Thalmann, M.Kruithof de Julio

## Abstract

Prostate cancer (PCa) is the second leading cause of cancer-associated death in men with therapy resistance acquisition to androgen deprivation treatment and metastasis progression. Understanding the mechanisms of tumor progression to metastatic stage is necessary for the design of therapeutic and prognostic schemes. The main objective of the current study is to determine, using transcriptomic and proteomic analyses on patient derived-xenograft models, whether differentially aggressive PCa tumors predispose their microenvironment (stroma) to a metastatic gene expression pattern, and how this information could be applied in prognostics. Transcriptomic profiling (RNA Sequencing) was performed on PCa PDX models representing different disease stages; BM18 (androgen dependent bone metastasis) and LAPC9 (androgen independent bone metastasis). Using organism-specific reference databases, the human-specific transcriptome, representing the tumor, was identified and separated from the mouse-specific transcriptome (representing the contributing stroma counterpart) from the same PDX tumor samples. To identify proteome changes in the tumor (human) versus the stroma (mouse), we performed human and mouse cell separation using the MACS mouse depletion sorting kit, and subjected protein lysates to quantitative TMT labeling and mass spectrometry. We show that tenascin C is one of the most abundant stromal genes in bone metastasis PCa PDXs, is modulated by androgen levels *in vivo* and is highly expressed in castration resistant LAPC9 PDX compared to castration sensitive BM18 PDX. Tissue microarray of primary PCa samples (N=210) was used to evaluate the potential of TNC to act as a metastasis prognosis marker. Low number of TNC-positive cells were associated with statistically significant clinical progression to local recurrence or metastasis, compared to high TNC-positive group. Our data showed that metastatic PCa PDXs that differ in androgen sensitivity trigger a differential stroma response suggesting that stroma was influenced by tumor cues. Selected stromal markers of osteoblastic PCa induced bone metastases, were induced in the microenvironment of the host organism in metastatic xenografts, although implanted in a non-bone site, indicating a conserved mechanism of tumor cells to induce a stromal pre-metastatic signature with high potential prognostic or diagnostic value.

## INTRODUCTION

Bone metastases are detected in 10% of patients already at the initial diagnosis of PCa or will develop in 20-30% of the patients subjected to radical prostatectomy and androgen deprivation therapy and will progress to advanced disease called castration resistant prostate cancer [1]. Metastases are established when disseminated cancer cells colonize a secondary organ site. An important component of tumor growth is the supportive stroma: the extracellular matrix (ECM) and the non-tumoral cells of the matrix microenvironment (e.g. endothelial cells, smooth muscle cells, cancer-associated fibroblasts). Upon interaction of stroma compartment and tumor cells, the stroma responds by secretion of growth factors, proteases and chemokines, thereby facilitating the remodeling of the ECM and thus tumor cell migration and invasion [2]. Therefore, tumor cell establishment requires an abnormal microenvironment. It is unclear whether the stroma is modulated by the tumor cells or by intrinsic gene expression alterations. Understanding the mechanisms of tumor progression to metastatic stage is necessary for the design of therapeutic and prognostic schemes.

The bone microenvironment is favorable for growth of PCa as well as breast cancer, indicated by the high frequency of bone metastasis in these tumors. Studies have shown that cancer cell growth competes for the hematopoietic niche in the bone marrow with the normal residing stem cells [3], and depending on the cancer cell phenotype, this may lead to either osteoblastic or osteolytic lesions. Stroma signature of osteolytic PCa cells (PC-3) xenografted intraosseously in immunocompomised mice induce a vascular/axon guidance signature [4]. Stroma signature of osteoblastic lesions from human VCap and C4-2B PCa cell lines, indicated an enrichment of hematopoietic and prostate epithelial stem cell niche. A curated prostate-specific bone metastasis signature (Ob-BMST) implicated seven highly upregulated genes (*Aspn*, *Pdgrfb*, *Postn*, *Sparcl1*, *Mcam*, *Fscn1* and *Pmepa1)* [5] among which *Postn* and *Fscn1* are bone specific. Furthermore, *Aspn* and *Postn* expression is also increased in primary PCa cases [5] indicative of osteomimicry processes. The induction of osteoblastic genes in the stroma of primary tumors (PCa and Breast), such as osteopontin and osteocalcin, has been suggested as a mechanism termed osteomimicry [6] to explain why the bone microenvironment is the preferential metastasis site. High stromal differences between benign, indolent and lethal PCa, combined with enrichment of bone remodeling genes in high Gleason score cases [7], suggest that stroma is an active player in PCa. During androgen deprivation, androgen dependent epithelial cells will undergo apoptosis, while the supporting stroma is largely maintained or replaces the necrotic tissue areas [8]. Stromal cells do express Androgen Receptor (AR) and have active downstream signaling, while absence of stromal AR expression is used as a prognostic factor of disease progression [9]. Furthermore, AR binds to different genomic sites in prostate fibroblasts compared to epithelium [10] and to cancer associated fibroblasts (CAFs) [11], indicating different roles of AR in epithelial or stroma cellular contexts. Prostate CAFs have tumor promoting effects on marginally tumorigenic cells (LNCaP), irreversibly alter their phenotype and influence progression to androgen independence and metastasis [12, 13].

In this study, we investigated whether metastatic PCa patient derived xenograft models (PDXs) that differ in androgen sensitivity are triggering a differential stroma response. To elucidate the mechanisms of stroma contribution to tumor growth later on, we determined the unique gene expression profile of stroma compared to the tumor compartment, the proteome changes of tumor versus stroma, and identified tumor-stroma interactions (Tenascin C-Integrin a2) of potential prognostic value in primary PCa.

## RESULTS

### Simultaneous transcriptome analysis of human and murine signatures in PDXs can distinguish androgen dependent expression changes in tumor and host-derived stroma

We analyzed the transcriptome of bulk PDX tumors grown subcutaneously in immunocompromised murine hosts by Next-Generation RNA Sequencing (RNASeq). Bone metastasis BM18 and LAPC9 PDXs were used in three different states; intact, post castration (day8 LAPCa9, day14 BM18) and androgen replacement (24hrs) (**Fig.1A**). Tumor growth kinetic revealed the androgen dependent phenotype of BM18, which regressed completely in 2 weeks post castration (**Fig.1B**) and the androgen independent phenotype of LAPC9 PDX tumors, which grew exponentially even after castration (**Fig.1C**) thus, confirming the differential aggressiveness of the two models. Reduction of epithelial glands and proliferating Ki67+ cells in the BM18 castrated conditions (**Fig.1D**) was in contrast to the LAPC9 tumors (**Fig.1D**) which were morphologically indistinguishable among intact and castrated hosts. Bulk tumor tissue which contain human tumor cells and mouse infiltrating stroma cells, were simultaneously analyzed from the same samples by RNASeq. To distinguish the transcriptome of the different organisms, the mouse and human reads were separated by alignment to a mouse and a human reference genome, respectively. Principal component analysis (PCA) of the human (tumor) 500 most variable genes showed that both castrated and replaced groups have altered expression profiles among each other and compared to the intact tumors. This was the case for the BM18 (**Fig.2A**) and the LAPC9 human transcriptome (**Fig.2B**). The response to short term androgen replacement showed a larger degree of variability in the BM18 (**Fig.2A**). However, expression levels of direct AR target genes (*KLK3, NKX3.1, FKBP5*) identified by the RNASeq, confirmed that androgen levels have affected the activation of androgen receptor signaling in both BM18 (**Fig.2C**) and LAPC9 (**Fig.2D**, *KLK3, NKX3.1*).

**Figure 1.**
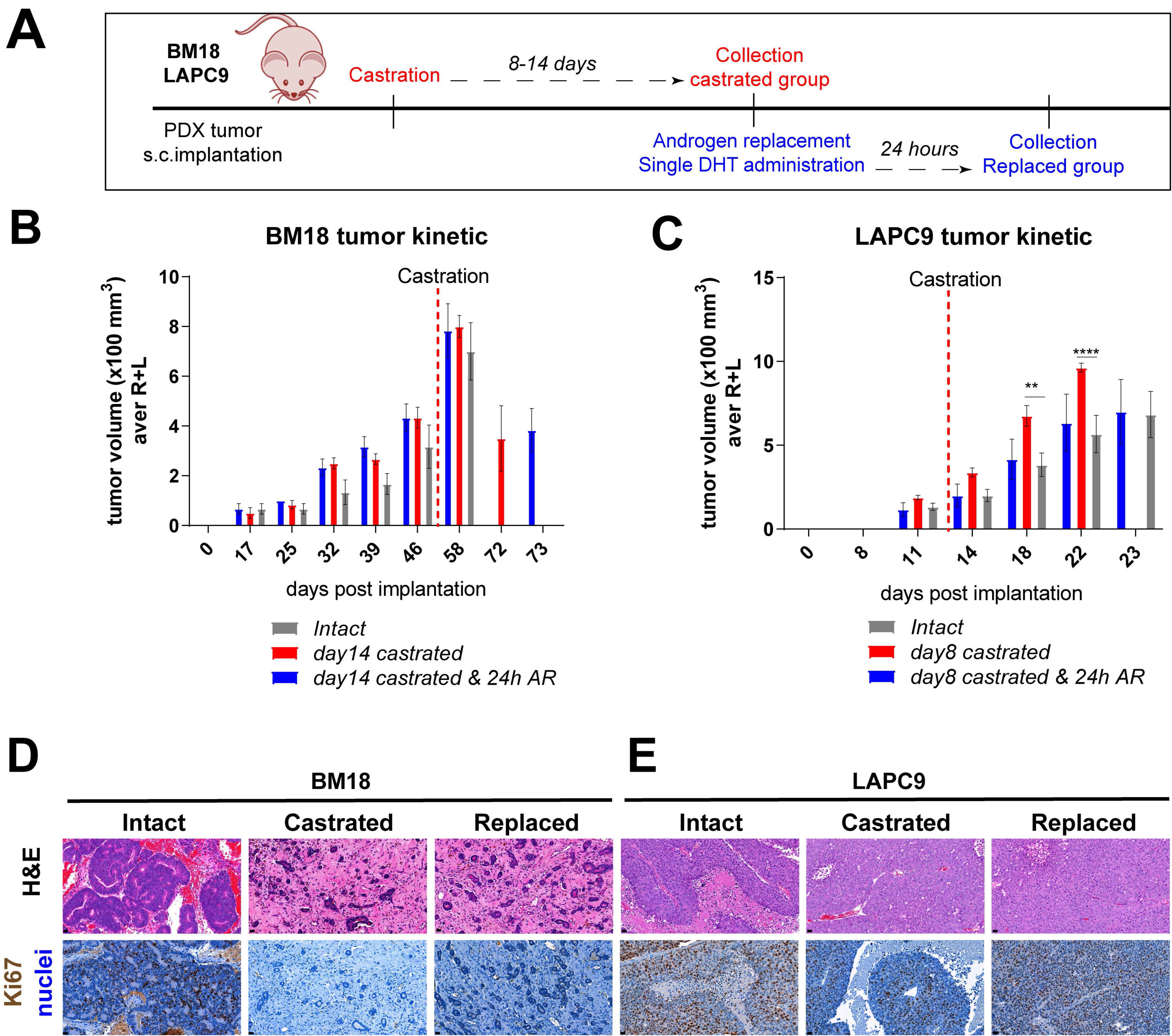
*In vivo* tumor growth properties of androgen-dependent BM18 versus androgen–independent PDX models. **A.** Scheme of *in vivo* BM18 and LAPC9 experiments, including timeline of castration, androgen replacement (single DHT testosterone administration) and collection of material for transcriptomic analysis. **B.** BM18 PDX tumor growth progression in time. Groups; 1.Intact tumors (collected at max size, N=3), 2.Castrated (day14, N=4), 3.Castrated followed by Testosterone re-administration (Castrated-Testosterone) (day15 since castration, 24hrs since AR, N=3). **C.** LAPC9 PDX tumor growth progression in time. Groups; 1.Intact tumors (collected at max size, N=3), 2.Castrated (day8, N=4), 3.Castrated followed by Testosterone re-administration (Castrated-Testosterone) (day9 since castration, 24hrs since AR, N=3). Tumor scoring was performed weekly by routine palpation; values represent average calculation of the tumors of all animals per group (considering 2 tumors, left L and right R of each animal). Error bars represent SEM, is calculated considering No of animals for each time point. Ordinary two-way ANOVA with Tukey`s multiple comparison correction was performed, p ˂ 0.01 (**), p ˂ 0.0001 (****). **D.** Histological morphology of BM18 and LAPC9 (from intact, castrated and androgen replaced hosts), as assessed by Hematoxylin and Eosin staining (H&E, top). Scale bars 20um, and proliferation marker Ki67 protein expression (bottom panel). Scale bars 20um.

**Figure 2.**
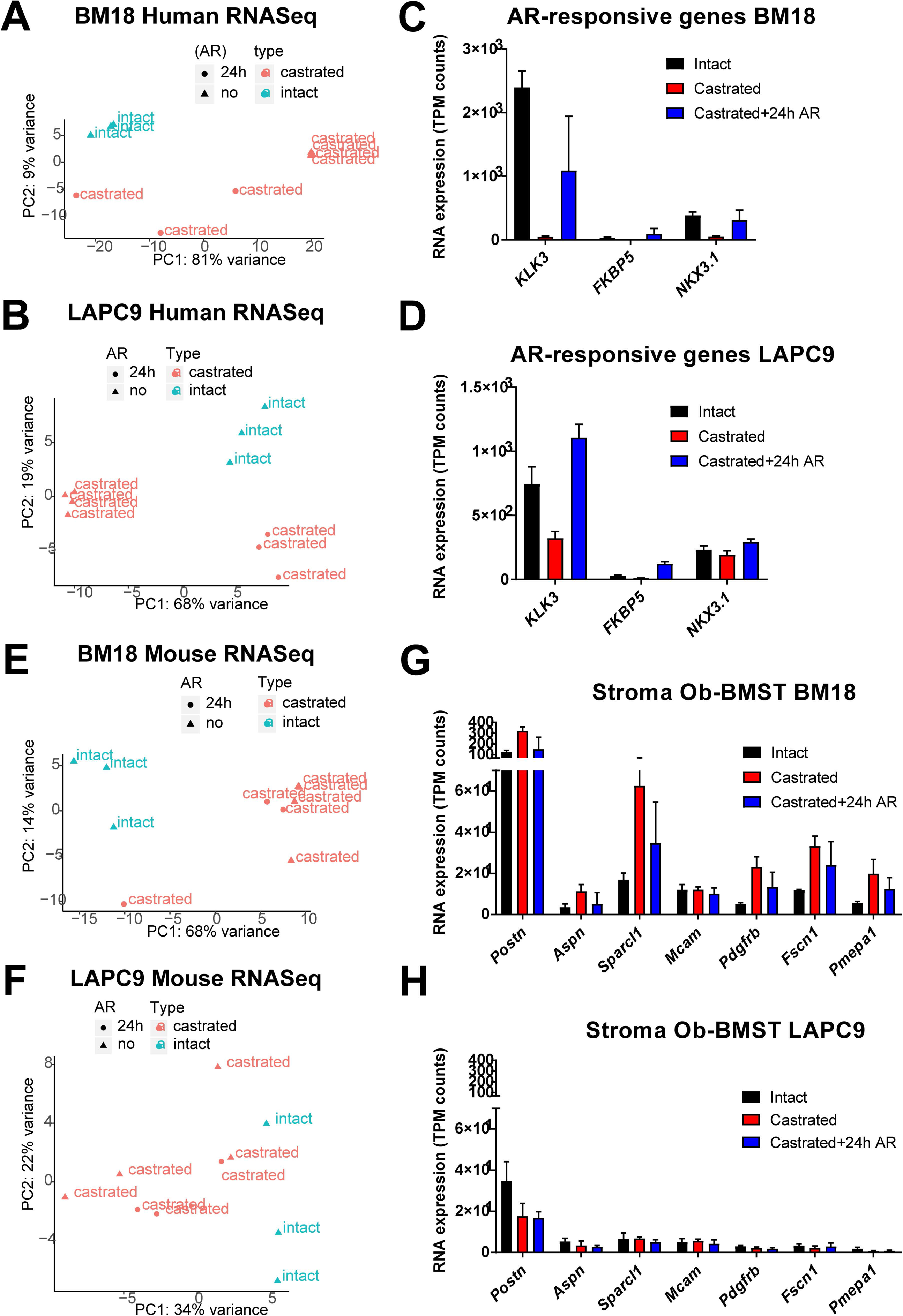
Separation of human (tumor) and mouse (stroma) transcriptome of BM18 and LAPC9 tumors. **A-B.** Principal component analysis plot of the gene expression of the 500 most variable genes on all samples, BM18 human transcripts (A), LAPC9 human (B) at intact, castrated, replaced (castrated+ 24h AR) conditions. **C-D.** Expression values of AR direct target genes as detected by RNASeq (TPM counts) in the BM18 (C) or LAPC9 (D) tumors as confirmation of effective repression of AR downstream signaling by castration. Intact (N=3), castrated (N=4), replaced (N=3). **E-F.** Principal component analysis plot of the gene expression of the 500 most variable genes on all samples, BM18 mouse (E) and LAPC9 mouse (F) at intact, castrated, replaced (castrated+ 24h AR) conditions. **G-H.** Expression values of Ob-BMST seven-upregulated stroma signature genes as detected by RNASeq (TPM normalized counts) in the mouse transcriptome of BM18 (E) or LAPC9 (F) tumors.

PCA analysis of the BM18 mouse (stroma) transcriptome indicated that the majority of castrated samples (with and without 24 h androgen replacement) diverge from the intact tumor (**Fig.2E**). LAPC9 mouse (stroma) transcriptome instead did not show specific clustering within or between sample groups when plotting the top 500 most variably expressed genes (**Fig.2F**). The Ob-BMST signature all seven genes (*Aspn*, *Pdgrfb*, *Postn*, *Aspn*, *Sparcl1*, *Mcam*, *Fscn1* and *Pmepa1),* which were upregulated in bone stroma as previously identified [5], were indeed expressed in both BM18 and LAPC9 PDXs with *Pdgrfb*, *Postn*, *Aspn*, and Sparcl1 specifically in the mouse RNASeq. Some of these genes were differentially expressed upon castration in the BM18 (**Fig.2G**) but not in the LAPC9 (**Fig.2H**).

### Proteomic analysis provides functional information over the identified human/mouse-specific transcriptome

To study the proteome of the tumor versus the stroma, human and mouse cell fractions were isolated by MACS Mouse Depletion method from tumor sample preparations; BM18 and LAPC9 each at intact, castrated, replaced state. Protein lysates of either mouse or human origin (single replicate from pool of N=3-4 biological replicates per condition) were subjected to an in-solution tryptic digest following TMT-labeling of the resulting peptides and their mass spectrometric analysis **(Fig.3A)**.

**Figure 3.**
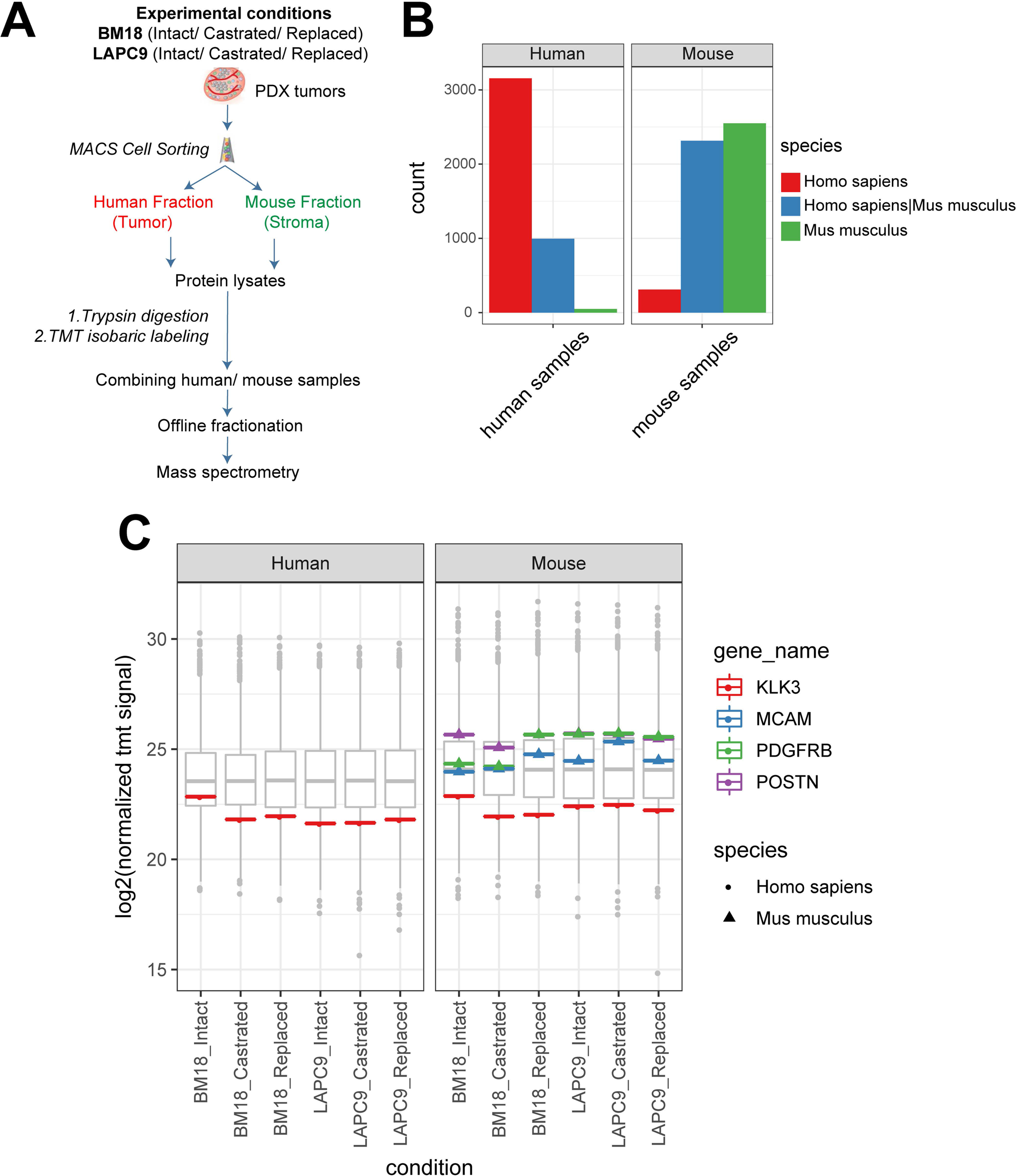
Proteomic analysis of human (tumor) versus mouse (stroma) of BM18 and LAPC9 tumors. **A.** Experimental separation of human from mouse cell suspensions from fresh tumor isolation by MACS Mouse depletion sorting. Cell fractions from intact/replaced (N=3 each), castrated (N=4) biological replicates were pooled into a single replicate (N=1) to achieve adequate cell number for proteomic analysis (1×10^6 cells). Protein lysates from the different fractions of BM18/LAPC9 (intact, castrated, replaced) were subjected to TMT labelling (all mouse or all human samples were multiple–xed in one TMT experiment each) followed by mass spectrometry. **B.** Detected peptides from human and mouse fractions were searched against a combined human and mouse protein database. Number of species specific or shared proteins is indicated in different colors. **C.** KLK3 (PSA) protein levels (log2 normalized tmt signal sum values) in human cell isolations (left) and in mouse cell isolations (right), and the protein sequence was predicted as human-specific (spheres indicate Homo Sapiens sequence). Seven-up Ob-BMST signature markers POSTN, PDGFRB and MCAM protein levels were absent in human cell isolations (left), present in mouse cell isolations (right), while all the protein sequences were mouse-specific (triangles indicate Mus Musculus sequences).

In addition to the initial experimental separation of the protein lysates, we further explored the species homologs of the identified proteins by computational analysis using a combined human and mouse protein sequence database. We identified 4198 proteins in the sample that has been enriched for human cells. 3154 thereof were human-specific proteins, with 996 revealing a high homology shared among human and mouse, and only a fraction of 48 mouse-specific peptides. (**Fig.3B**, left plot). For samples enriched in mouse cells we identified in total 5192 proteins; thereof 2486 mouse-specific proteins, 2379 shared homologs, and 247 human-specific (**Fig.3B**, right plot). We searched for prostate specific markers such as KLK3, a prostate specific antigen, which is secreted by luminal cells. In the proteomic data the human-specificity was confirmed, and the secreted protein was found also in the mouse fraction (**Fig.3C**). To further ensure that the proteomic data were indeed identifying real stromal specific candidates, we searched specifically for the seven-gene Ob-BMST signature found also to be expressed in both BM18, LAPC9. POSTN, PDGFRB and MCAM (**Fig.3C**) were indeed detected at the protein level, thus might have a functional role, and were found exclusively in the mouse fraction (**Fig.3C**, right plot) and hybridizing with mouse-specific sequence (**Fig.3C**, triangle indicates Mus Musculus species specificity).

### Differential expression analysis reveals androgen dependent stromal gene modulation in androgen independent PDX model

We demonstrated that the human (tumor) as well as the mouse (stroma) transcriptomes follow androgen dependent transcriptomic changes in the BM18 groups (Intact vs castrated vs replaced) (**Fig.2A,E**). To identify the top most significant AR regulated stromal genes we performed differential expression analysis of BM18 tumors (**Fig.4A**) from castrated hosts and compared it to BM18 intact (the replaced tumors were not included here due to higher variability). Of the top most variable genes, 50 were highly upregulated in BM18 tumors (Z score >1) and downregulated upon castration (**Fig.4A**). Differential expression analysis of LAPC9 tumors from castrated/replaced tumors, versus intact tumors revealed the top most differentially regulated genes; 27 most upregulated genes in the intact, which were downregulated in castrated groups (**Fig.4B**). Among the 50 mouse genes that were highly upregulated in the intact BM18, and significantly modulated by castration, were 23 genes implicated in cell cycle/mitosis, 10 implicated in ECM, 3 related to spermatogenesis/hormone regulation, according to Gene Ontology terms (**Fig.4C**). Two of these genes *Tnc* and *Crabp1*, were also detected in the proteomic data (**Fig.4C**, highlighted in bold) and in both PDXs (**Fig.4C-D**, highlighted in red). Among the 27 mouse genes that were highly upregulated in the intact LAPC9, and significantly modulated by castration, 7 genes implicated in ECM/cell adhesion/ smooth muscle function and 14 implicated in non-smooth muscle function and metabolism based on Gene Ontology terms (**Fig.4D**). In the LAPC9 proteomic data we detected 14 genes out of the 27 to be expressed in the mouse fractions (**Fig.4D**, bold), indicative of potential functional value. Of interest in potentially mediating tumor stroma extracellular interactions are; Neural adhesion protein (*CD56*), implicated in cell-cell adhesion and migration by homotypic signaling, as well as Tenascin C (*Tnc*), an extracellular protein which is found abundant in reactive stroma of various cancer types, yet not expressed in normal stroma. Both genes were expressed at the protein level, exclusively in the mouse compartment of the BM18 and LAPC9, at all states (Intact, castrated, replaced). Furthermore, Tnc was detected in both BM18, LAPC9 at the transcriptional and proteomic level, and is reactivated at 24hours of androgen replacement (**Fig.4B**), indicative of AR-direct target gene modulation.

**Figure 4.**
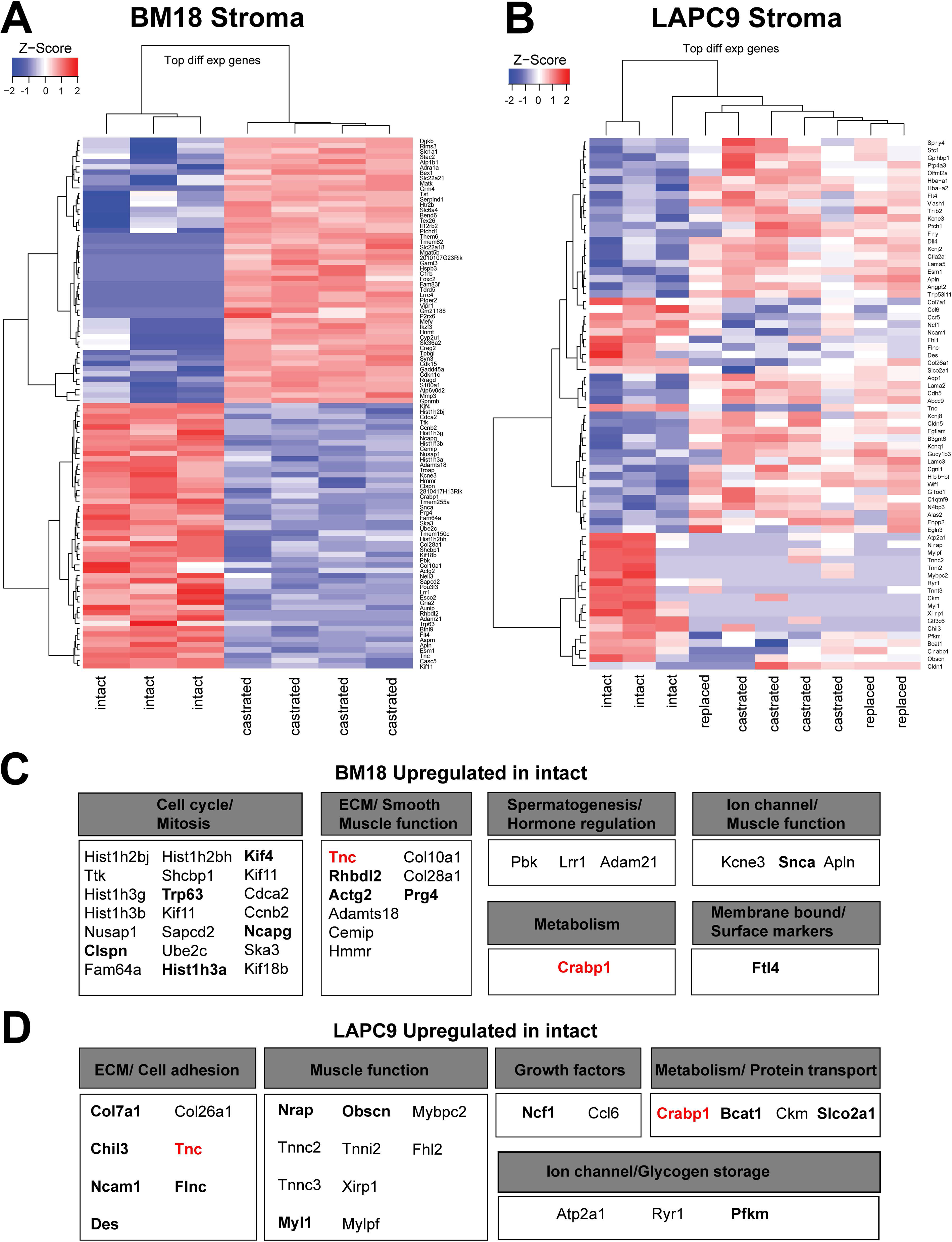
Differential expression analysis of transcriptome indicates different expression profiles of stromal genes as response to androgen deprivation. **A.** Heatmap represents differential expression analysis of the most variable genes from mouse transcriptome of BM18 castrated compared to BM18 intact tumors. Genes modulated by androgen deprivation due to castration in up/downregulation compared to intact tumors are indicated in red or blue color, respectively. **B.** Heatmap represents Z-score of differential expression analysis of most variable genes in mouse transcriptome of LAPC9 castrated (with and without androgen replacement) compared to LAPC9 intact tumors. **C.** Description of mouse genes found upregulated in BM18 intact tumors and the biological processes they are involved in according to GO terms. **D.** Description of mouse genes found upregulated in LAPC9 intact tumors and the biological processes they are involved in, according to GO terms.

### Cross comparison of stromal transcriptome among different PDXs identifies Tenascin C upregulated in castrated LAPC9 androgen independent model

To assess the similarity between the stromal transcriptome of the androgen independent LAPC9 and the BM18, differential expression analysis was performed. In a panel of 50 top most variable genes comparing the tumors at their intact condition, we identified several genes that follow the same pattern of modulation in intact tumors (**Fig.5A**) and in castrated tumors (**Fig.5B**). Of interest were the ECM related genes downregulated in LAPC9 versus BM18; the FGF receptor (*Fgfr4*), elastin microfibril interface (*Emilin3*) and upregulated Collagen type 2 chain a1 (*Col2a1*). The differential expression of LAPC9 castrated versus BM18 castrated, highlights genes that were identified in the analysis among LAPC9 castrated, replaced versus LAPC9 intact such as Apelin (*Apln*), *Col2a1* and Tenascin C (*Tnc*). Given that genes activated in castrated state might be indicative of androgen resistance mechanism activation, we postulated that genes upregulated in the androgen resistant LAPC9 over the androgen dependent BM18 might be relevant for understanding the aggressive phenotype of LAPC9 and therefore of advanced metastatic phenotype of similar tumors. Tenascin is an ECM protein which is produced at the (myo)fibroblasts that is virtually absent in normal stroma in prostate and other tissues, and has been associated with cancerous reactive stroma response in different cancers. We interrogated the expression of *Tnc* in the RNASeq data and found that it was highly upregulated in LAPC9 compared to BM18 both in intact (logFC 4.23, p value ˂0.001) and among the castrated conditions (logFC 6.9, p value ˂0.001) (**Fig.5C**). However, in both models *Tnc* levels significantly decrease upon castration (BM18, p value ˂0.001), LAPC9, p value ˂0.05), indicating potentially AR-mediated regulation of *Tnc* expression.

**Figure 5.**
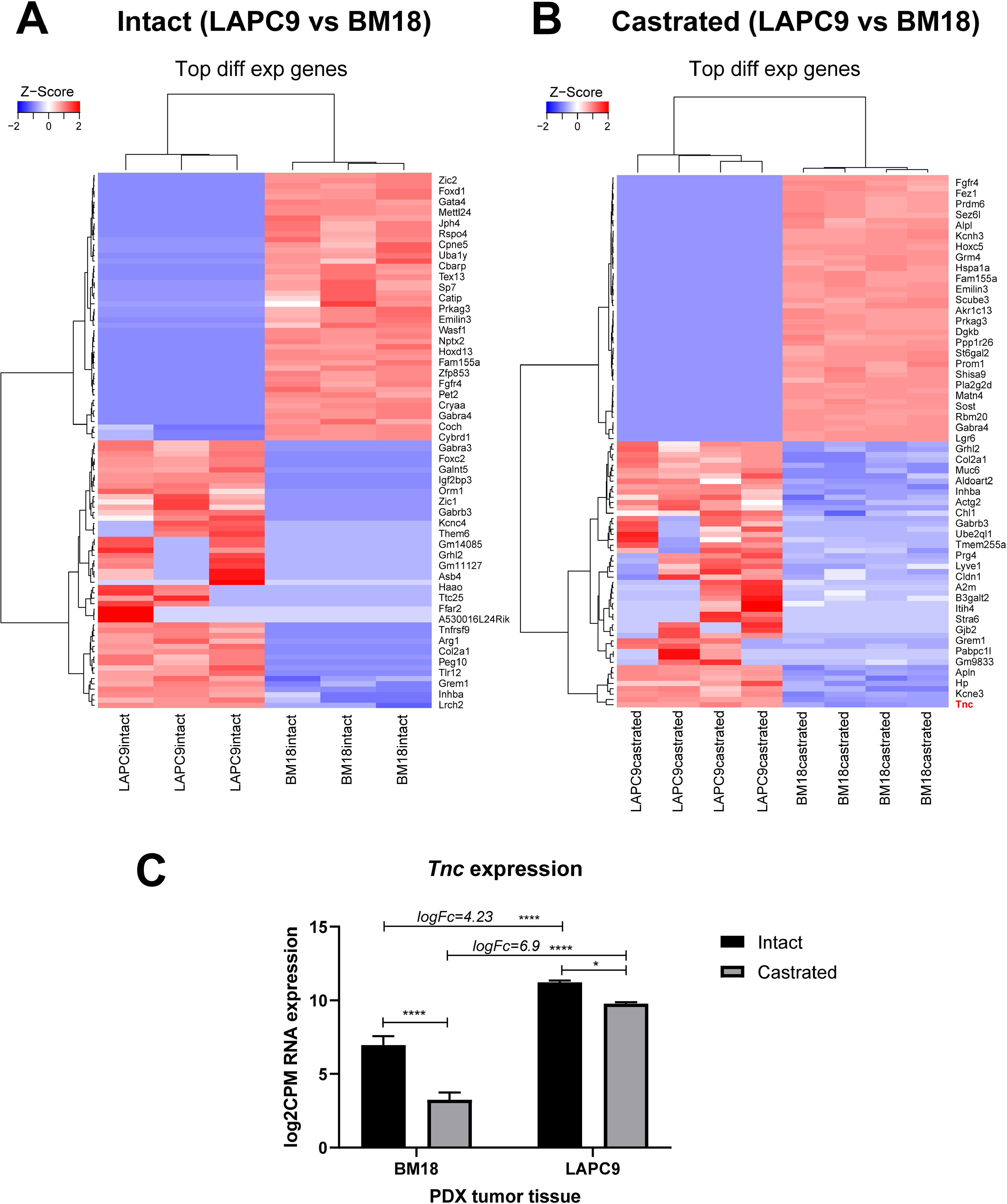
Crosscomparison of LAPC9 versus BM18 suggests stromal gene Tenascin C expression to be associated with advanced PCa and regulated by androgen levels. **A.** Heatmap represents differential expression analysis of the top100 most variable genes from mouse transcriptome of LAPC9 intact tumors compared to BM18 intact tumors and **B.** of LAPC9 castrated tumors compared to BM18 castrated tumors. A subset of genes in LAPC9 samples have zero counts, leading to same z score, while the same genes are highly expressed in BM18 samples. **C.** *Tnc* RNA expression (log2CMP counts) in stroma transcriptome. LogFc enrichment of Tnc in LAPC9 over BM18 is indicated. Ordinary two-way ANOVA with Tukey`s multiple comparison correction was performed, p ˂ 0.05 (*), p ˂ 0.0001 (****).

### Protein expression of Tenascin and its interaction partners

To assess whether the transcriptomic changes of *Tnc* in the PDX models, corresponds to functional protein and thus a relevant role in bone metastatic PCa, we used proteomic analysis. Mass spectrometry analysis of the human and mouse fractions indicated that Tnc protein was expressed specifically in the mouse (stromal) fractions in BM18 and LAPC9 (**Fig.6A**). The isoform Tenascin X was also expressed at the protein level (**Fig.6B**). The interaction network of Tnc mouse protein Tnc is based on experimental observations and prediction tools (STRING), consists of laminins (Lamc1, Lamb2), fibronectin (Fb1), Integrins (Itga2, a7, a8, a9) and proteoglycans (Bcan, Vcan) (**Fig.6C**). The human interactome is less characterized, yet most of the interactome is conserved; laminins (LAMC1, LAMB2), proteoglycans (NCAN, ACAN), and others such as IL-8, BMP4, ALB, SDC4 (**Fig.6D**). However, integrin interaction binding partners in a human setting has not been confirmed. Given the importance of integrins for cell adhesion and migration known to be found in mesenchymal/stromal and epithelial tumor cells, we focused on the expression of human- and mouse-derived integrins. The *ITGA9*, *ITGA6*, *ITGA2*, were all found to be expressed in both the RNASeq and proteomic data (**Fig.6E**, ITGA6 respectively), however only *ITGA2* protein was specifically found in the human counterpart and not overlapping with the mouse stroma (**Fig.6F**). Tumor *ITGA2* and stromal *Tnc* is therefore a potential molecular interaction, part of the dual communication among tumor and its microenvironment with promising implications for therapeutic targeting.

**Figure 6.**
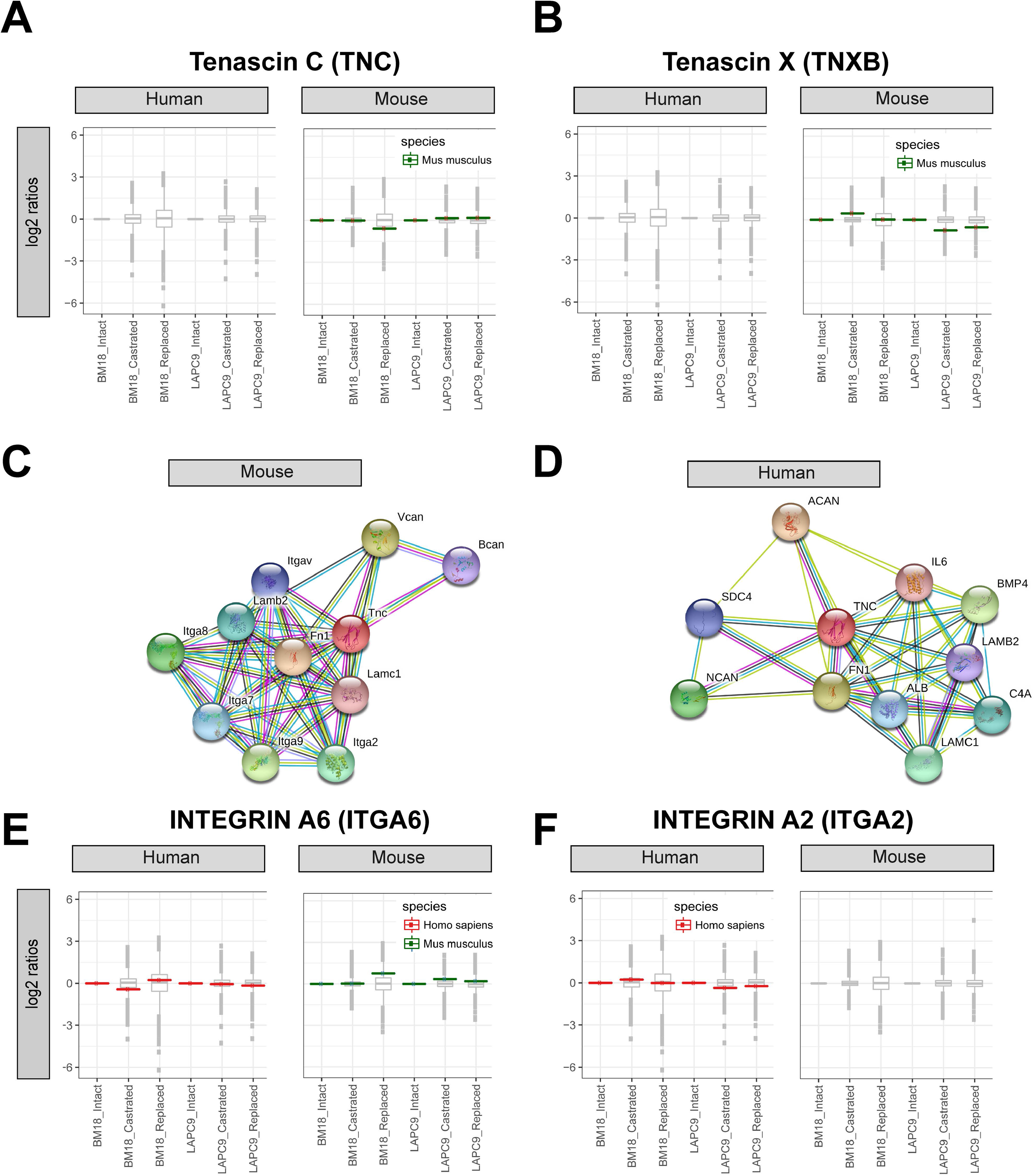
Tenascin C and its predicted interaction partners analysed by mass spectrometry. **A.** Tenascin C (TNC) and **B.** alternative isoform Tenascin X (TNXB) protein relative abundance (log2 ratios, single replicate per sample from a pool of n=3-4) in human cell isolations (left) and present in mouse cell isolations (right). The vsn corrected tmt reporter ion signals were normalized by the intact conditions of either BM18 or LAPC9. The protein sequences were predicted as mouse-specific (green). **C.** Protein interaction network of mouse TNC protein based on STRING association network (https://string-db.org/). **D.** Protein interaction network of human TNC protein based on STRING association network https://string-db.org/. **E.** TNC binding partner Integrin A6 (ITGA6) was detected by mass spectrometry in both the human and mouse protein lysates and matching the organism-specific protein sequence based on bioinformatics analysis (red for human, green for mouse). **F.** TNC binding partner Integrin A2 (ITGA2) was detected by mass spectrometry specifically in the human protein lysates and was matching the human-specific protein sequence.

### Stromal Tenascin expression as a prognostic factor of disease progression in high risk PCa

The detection of key mouse stromal genes in PCa PDXs gives the opportunity to evaluate the role and potential prognostic value of the human orthologs of these stromal genes. To validate the localisation and stromal specificity of TNC protein expression we performed immunohistochemistry on the LAPC9 and BM18 PDX and primary PCa tissue sections. TNC is localised in the extracellular space (**Fig.7A**, *primary cases*) and in morphologically myofibroblast cells (**Fig.7A**, BM18, LAPC9) in proximity to the vessels (**Fig.7A**, *LAPC9*). Next we evaluated TNC expression in a tissue microarray of 210 primary prostate tissues part of the European Multicenter High Risk Prostate Cancer Clinical and Translational research group (EMPaCT) [14–16] (**Fig.7B-G**). Based on the preoperative clinical parameters of the TMA patient cases (**Table 1**) and the D’Amico classification system [17], they represent intermediate (clinical T2b or Gleason N=7 and PSA >10 and ˂=20) and high risk (clinical T2c-3a or GS=8 and PSA>=20) PCa. Number of TNC-positive cells (**Fig.7B**) were quantified and averaged for all cores (four cores per patient case) in an automated way including, tissue selection, core annotation, and equal staining parameters set. To investigate the association between the number of TNC-positive cells and patient survival or disease progression, we calculated the optimal cut-point for the number of TNC-positive cells by estimation of maximally selected rank statistics [18]. Association between TNC-expressing cells and pTStage indicated that the majority of cases cluster towards stages 3a and 3b (**Fig.7C**). Multiple comparison test among all groups showed no statistically significant association between TNC expression and pathological stage (**Table 2**, p>0.05). Overall survival probability between two patient groups with respectively high and low number of TNC-positive cells was indifferent (p=0.29, Log-rank test) (**Fig.7D**). We focused on the probability of TNC expression in primary tumors to be a deterministic factor for clinical progression to local or metastasis recurrence. Clinical progression probability was higher in the TNC-low group compared the TNC-high group (p=0.04, *, Log-rank test) (**Fig.7E**). Next, we examined clinical progression in patients with pT Stage ≥3 (groups 3a, 3b and 4). The high T Stage cases did separate into two groups based on TNC expression, with TNC-low expressing group exhibiting earlier clinical progression (local or metastatic recurrence, p=0.013, *, Log-rank test) (**Fig.7F**). PSA progression probability in patients with pT Stage ≥3 indicated an association trend of TNC-low group with earlier biochemical relapse events (p=0.07, Log-rank test) (**Fig.7G**). Similarly, TNC-low group correlated with higher probability for PSA progression after radical prostatectomy among cases with carcinoma-containing (positive) surgical margins (**Sup.Fig.2A**, p=0.031, *, Log-rank test) or positive lymph nodes (**Sup.Fig.2B**, p=0.092, Log-rank test). Low number of TNC expressing cells coincides with poor prognosis in terms of metastasis progression, similarly to its downregulation upon castration in the bone metastasis PDXs (**Fig. 4B**) based on the RNASeq analysis.

**Figure 7.**
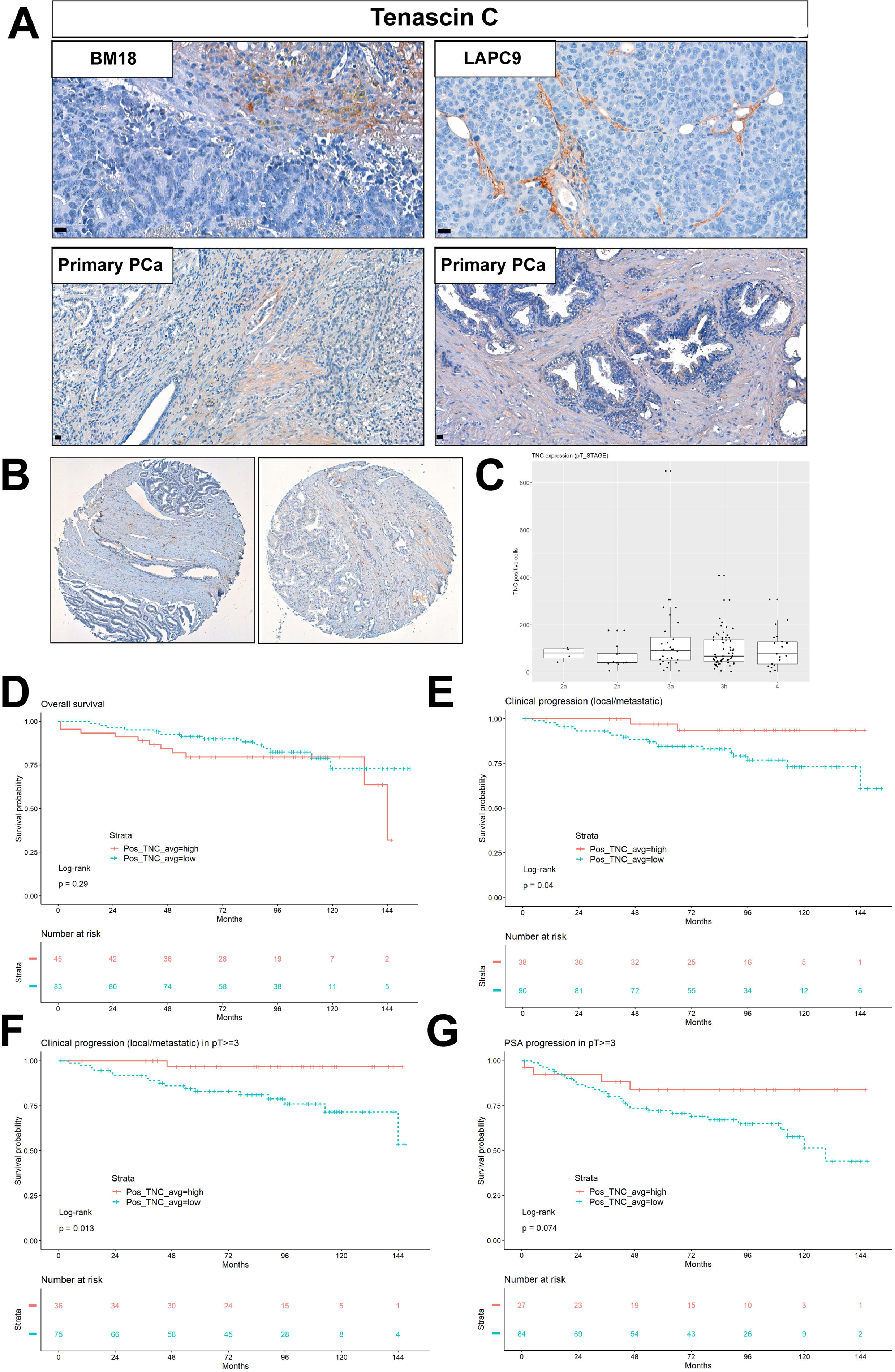
TNC protein expression in PDXs and primary, high risk, PCa TMA cases. **A.** Validation of protein expression and stromal specific of Tnc by immunohistochemistry in BM18 intact tumors, LAPC9 intact tumor, and primary PCa cases. **B.** Representative cases of TNC staining on primary PCa EMPACT TMA. **C.** TNC expression levels in terms of No positive cells in p T Stage classification. Statistical multiple comparison test Wilcoxon rank sum test was performed. p>0.05 **D.** Overall survival probability in patient groups of TNC-high and TNC-low (No of positive, TNC expressing cells) (p=0.29, ns). Average value represents the mean of four cores per patient case. **E.** Clinical progression to local recurrence or metastasis probability in patient groups of TNC-high and TNC-low expression (p=0.04, *˂0.05). **F.** Clinical progression to local recurrence or metastasis probability among patients of pTStages 3a,3b and 4 based on TNC-high and TNC-low expression (p=0.013, *˂0.05). **G.** PSA progression probability among patients of pTStages 3a,3b and 4 based on TNC-high and TNC-low expression (p=0.074, ns).

**Table 1.** Clinical parameters of EMPACT TMA patient cases.

|  | **Age at surgery** | **PSA at surgery** | **PSA progression time (months)** | **Clinical progression time (months)** |
| --- | --- | --- | --- | --- |
| **Min** | 43 | 20 | 1 | 1 |
| **1st quartile** | 62 | 25.33 | 29.5 | 40.5 |
| **Median quartile** | 67 | 36.99 | 63.5 | 75.5 |
| **Mean quartile** | 66.18 | 50.56 | 63.47 | 70.89 |
| **3rd quartile** | 71 | 61.9 | 90 | 95.75 |
| **Max quartile** | 81 | 597 | 151 | 153 |

|  |  | **Pathological Staging (No of patient cases)** |  |  |  |  |
| --- | --- | --- | --- | --- | --- | --- |
|  |  | **2a** | **2b** | **3a** | **3b** | **4** |
| **PSA progression** | **Clinical Progression** |  |  |  |  |  |
| **no** | no | 6 | 15 | 37 | 63 | 26 |
|  | yes | 0 | 0 | 0 | 1 | 0 |
| **yes** | no | 0 | 5 | 7 | 11 | 7 |
|  | yes | 1 | 7 | 9 | 9 | 6 |

**Table 2.** Statistical test TNC expression in patient groups of different pT Stage classification in the high risk PCa TMA. Shapiro statistical test for pT Stage 3a, 3b, (p values ˂0.001, ****), and group 4 (p>0.05, non significant). Data not homogeneously distributed, therefore non parametric one-way ANOVA and multiple comparisons using Wilcoxon rank sum test among pT Stage groups (2a, 2b, 3a, 3b, 4) were performed. Adjusted p values indicated (p>0.05, ns).

|  | 2a | 2b | 3a | 3b |
| --- | --- | --- | --- | --- |
| 2b | 0.86 | - | - | - |
| 3a | 1.00 | 0.86 | - | - |
| 3b | 1.00 | 0.86 | 0.86 | - |
| 4 | 1.00 | 0.86 | 1.00 | 1.00 |

P value adjustment method: BH

## DISCUSSION

The role of microenvironment upon cancer formation, and progression to metastasis is supported by numerous studies [19, 20], however the current knowledge is not sufficient to reconstruct the chain events from primary to secondary tumor progression. Normal stroma microenvironment is considered to halt tumor formation, however after interaction with tumor cells, it also undergoes a certain “transformation” at the transcriptomic and even at the genetic level [21–24]. The processes by which PCa tumor cells affect stroma, and in turn stroma impacts primary PCa tumor growth or metastasis are complex and remain largely unclear.

We utilised bone metastasis PDX models, of different aggressiveness with regards to androgen dependency. *In vivo* PDX models grafted in immunocompromised mice, although lack the complexity of a complete immune system, represent stroma compartment (endothelial cells, smooth muscle cells, myofibroblasts, cancer associated fibroblasts). Due to subcutaneous growth of BM PCa PDXs, human stroma is replaced by mouse infiltrating stromal cells and vasculature [25, 26]. Mouse cell infiltration allows the discrimination of organism-specific transcripts; human-derived transcripts representing the tumor cells, mouse-derived transcripts representing the mouse stroma compartment. Using next generation RNASeq, MACS based human and mouse cell sorting, mass spectrometry and organism-specific reference databases we have identified the tumor-specific (human) from the stroma-specific (mouse) transcriptome and proteome of bone metastasis PCa PDXs. The dynamics of AR signalling in the stroma are best represented in an *in vivo* setting [27], therefore to specifically examine the stroma changes dictated by PCa cells, we subjected the PDXs in androgen and androgen-deprived conditions. By imposing this selection pressure we could identify androgen dependent gene expression patterns.

We demonstrated that the human (tumor) as well as the mouse (stroma) transcriptomes follow androgen dependent transcriptomic changes in the BM18 groups (Intact vs castrated vs replaced). Despite the androgen independent tumor growth of LAPC9, at the gene expression level the LAPC9 tumor cells do follow AR-responsive pattern (human transcriptome). However, principal component analysis showed that although castrated and replaced LAPC9 groups separate adequately based on the human transcriptome, they appear to have overall uniform stromal transcriptome.

We report that transcriptomic mechanisms linked to osteotropism were conserved in bone metastatic PDXs, even in non-bone environment and differential stroma gene expression is induced by different tumors, indicating tumor-specificity of stroma reactivity. The Ob-BMST signature all seven genes (*Aspn*, *Pdgrfb*, *Postn*, *Sparcl1*, *Mcam*, *Fscn1* and *Pmepa1)* which were upregulated in bone stroma previously identified [5], were indeed expressed in both BM18 and LAPC9 PDXs specifically in the mouse RNASeq and also expressed at the protein level as identified by mass spectrometry. The gene expression modulation of mouse stroma is ultimately an important evidence of the effect of tumor cells in their microenvironment, where they induce favourable conditions for their growth.

Differential expression analysis of LAPC9 stroma signature from intact, castrated and replaced hosts, highlighted the most significantly variable genes which were modulated by androgen levels, despite the androgen independent tumor growth phenotype. Focusing on the genes that were highly activated in intact, but strongly modulated by castration, we categorized these genes based on Gene Ontology terms. We found that LAPC9 stromal genes were ECM remodeling components, and genes involved in smooth muscle function or even in striated muscle function. Of interest are *CD56*, *Tnc*, *Flnc*. Among the BM18 most abundant stromal transcripts, were genes involved in cell cycle regulation and cell division. Interrogating the differences among the two models, we focused on the transcriptome of LAPC9 normalised versus the less aggressive, androgen dependent BM18. In particular, *Tnc* is expressed in both PDXs, higher in the LAPC9, yet downregulated upon castration suggesting a direct AR gene regulation. Differential expression analysis among both the PDXs after castration, indicated that *Tnc* is upregulated in LAPC9 than BM18, suggesting an association with disease aggressiveness. Genes which become upregulated in castrated conditions are likely to be linked to androgen resistance, thus we studied *Tnc* for potential role in metastasis progression.

TNC is an extracellular glycoprotein, absent in normal prostate and in postnatally silenced in most tissues. TNC is re-expressed in reactive stroma in human cancers and there is evidence of its expression in low grade tumors (Gleason 3) of human PCa [28] and possibly already activated at PIN stage [29, 30]. In particular, high molecular weight TNC isoforms are expressed in cancer due to alternative mRNA splicing [29]. We examined whether abundance of TNC-positive cells in primary PCa TMA can predict metastatic progression and overall survival (12 years follow-up after radical prostatectomy). High number of TNC-positive cells did not correlate with overall survival or histological grade in agreement with previous data [29]. PSA progression after radical prostatectomy was occurring earlier in TNC-low group compared to TNC-high group when high stage cases (pT ≥3), surgical margin-positive or lymph node-positive cases were investigated. In terms of clinical progression, the TNC-low group in the total number of cases and among the high stage (pT ≥3) cases showed worse prognosis in terms of local recurrence/ metastasis. This finding is in contrast to the study of Ni et al., showing that high levels of TNC is significantly linked to lymph node metastasis and clinical stage [31] but in agreement with another study which reported weak TNC expression in high grade PCa [30]. No low risk cases or metastasis tissues were used in our study, and we focused on TNC-producing cells not overall TNC expression in the matrix. Therefore we can only conclude that TNC is indeed expressed in intermediate and high risk primary PCa as assessed at preoperative diagnosis based on D’Amico criteria [17], and that high number of TNC-positive cells is inversely correlated with clinical progression.

More evidence point to the direction that TNC might be degraded upon local recurrence in lung cancer [32, 33], while high TNC is found in lymph and bone metastases sites [29], or even in certain types of bone metastasis [34]. In TMA of PCa bone metastasis, San Martin et al., demonstrated high TNC expression in trabeculae endosteum the site of osteoblastic metastasis, and yet low TNC expression in the adjacent bone marrow sites [34]. Osteoblastic PCa cell lines proliferate rapidly *in vitro* and adhere to TNC protein while osteolytic PC3 or lymph node derived PCa lines do not show this phenotype, suggesting an association of TNC with osteoblastic but not osteolytic metastases. One of the ligands of TNC highly upregulated in VCap cells was α9 integrin which binds directly TNC and modulate expression of collagen [34], providing evidence for TNC-integrins in human PCa. Our RNASeq data indicate also expression of α9 integrin along with α6 and α2, and based on the proteomic human-mouse separation we found integrin α2 to be the only human-specific and thus tumor-specific for the PDXs used in this study. Although the molecular mechanism among TNC-ITGA2 should be further characterised, evidence on correlation among α2 and α6 expression in primary PCa and bone metastasis occurrence has been previously reported [35].

The reactivation of TNC expression is relevant for reactive stroma regulation, while TNC downregulation might be relevant for recurrence or metastasis initiation, which remains to be further investigated. Indeed, TNC is known to have pleiotropic functions in different cellular contexts with both autocrine TNC expression in tumor cells and paracrine TNC from stroma in different stages of metastasis [36], however the cellular source of TNC in primary PCa was not addressed in our study. Our data demonstrate that androgens regulate stromal TNC expression, evident by reduced TNC expression upon castration (even in the castration resistant LAPC9) and immediate increased expression upon androgen replacement, thus TNC expression should be further evaluated in CRPC samples. Genomic amplification in TNC gene associates with highly aggressive neuroendocrine PCa occurrence [37]. In a multi-omics approach study, TNC protein was one of the panel of four markers detected in preoperative serum samples and collectively predict biochemical relapse events with high accuracy [38].

In summary, we have identified the stroma signature of bone metastatic PDXs and by analysing androgen dependent versus androgen-independent tumors, we could demonstrate that tumor-specific stroma gene expression changes. We could show that there are AR-regulated stromal genes modulated upon castration, even in the androgen independent for tumor growth, LAPC9 model. Osteoblastic bone metastasis stromal seven-gene signature was induced in the mouse-derived stroma compartment of BM18 and LAPC9, indicating conserved tumor mechanisms which can induce transcriptomic “transformation” of mouse infiltrating stroma (even in subcutaneous sites), to bone microenvironment-like stroma. The prognostic value of stroma signatures has been also demonstrated by another study utilising PDXs associated with metastasis prognosis from different lesions from a single PCa case, and demonstrated the strong predictability of 93-gene stroma signatures to metastasis phenotypes in different clinical cohorts [39]. We have identified androgen dependent Tenascin C expression in the stroma of PDX models, which is downregulated in the conditions mimicking aggressive disease (upon castration), similarly to the high clinical progression probability of low TNC group in the primary PCa TMA. The higher stromal *Tnc* mRNA levels in the aggressive LAPC9 compared to BM18 may suggest that it would be relevant to examine TNC mRNA and protein expression in human bone metastasis or ideally matched primary-metastasis cases in order to understand the kinetic of TNC in terms of disease progression. Given that TNC expression was found elevated from 0% in BPH stroma to 47% in tumor-associated stroma [24], its detection in circulation [38], its immunomodulatory role [40], indicates TNC as a promising drug target and disease determining factor. TNC clinical progression predictive value is performing best in earlier stage, low risk PCa, while our data show that in high risk PCa low number of TNC producing cells were associated with poor prognosis possibly due to changes in tissue remodeling and thus variable TNC levels. The regime that metastatic, stroma-specific molecular signature may be detectable in the PCa site either prior to or during metastasis, will most likely require not a single marker approach but a combination of biochemical, histological markers taking into consideration dual tumor-stroma interactions in order to provide prognostic tools for improved patient stratification after initial PCa diagnosis and preventive surveillance for metastasis risk.

**Supplementary Figure 1.**
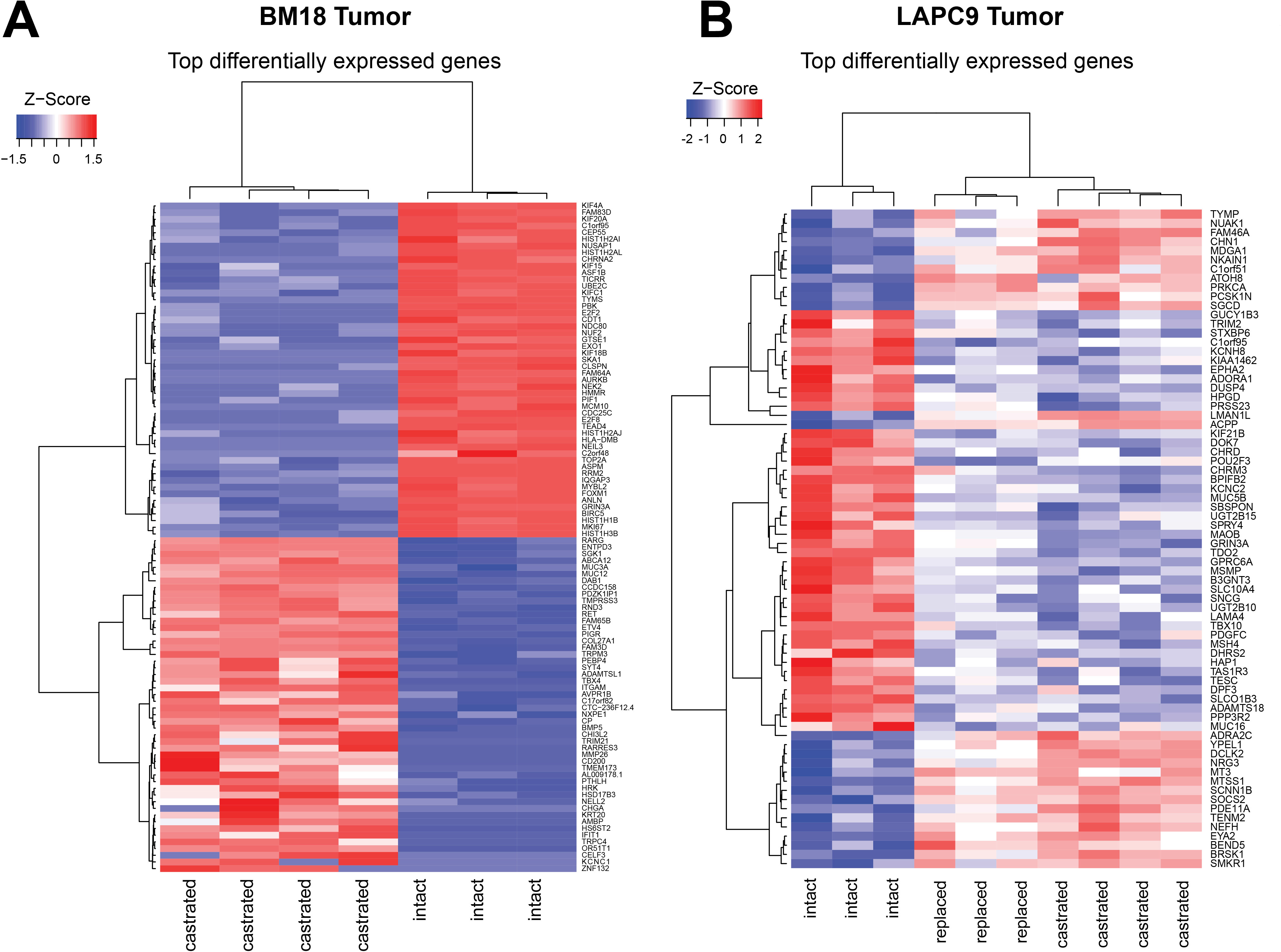
Human transcriptomic profile of BM18 and LAPC9 tumors. **A.** Heatmap represents differential expression analysis of most variable genes from human transcriptome of BM18 castrated compared to BM18 intact tumors. **B.** Heatmap represents differential expression analysis of most variable genes from human transcriptome of LAPC9 castrated (with and without androgen replacement) compared to LAPC9 intact tumors. Genes modulated under androgen deprivation conditions by up/downregulation compared to intact tumors are indicated in red or blue color, respectively.

**Supplementary Figure 2.**
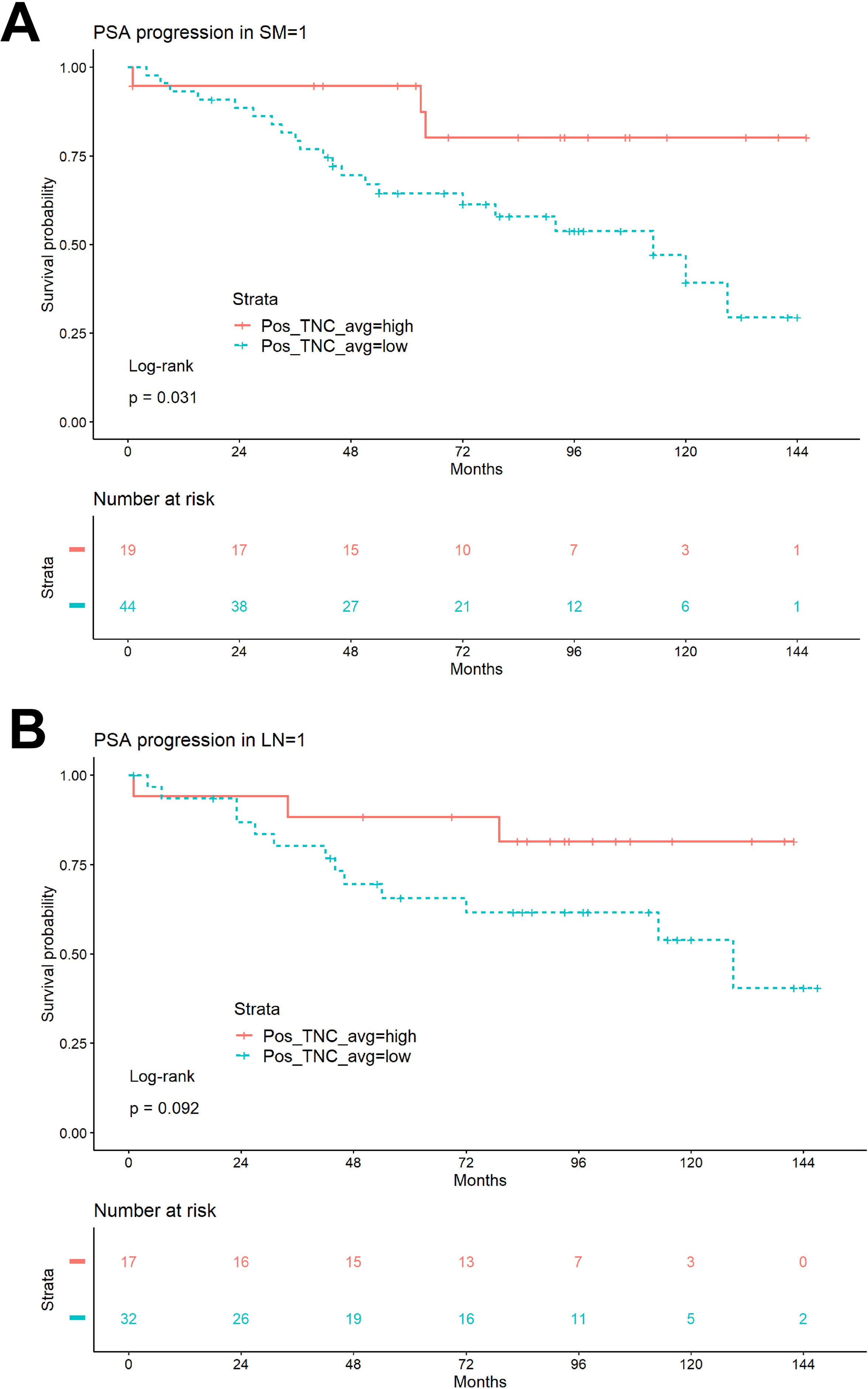
PSA progression in cases with positive surgical margins or lymph node status. **A.** PSA progression probability of patient cases with positive surgical margins presence (SM=1) according to TNC-high and TNC-low expression (p=0.031, *˂0.05). **B.** PSA progression probability of patient cases with positive lymph node (LN=1) status according to TNC-high and TNC-low expression (p=0.092, ns).

## MATERIALS AND METHODS

### Tumor sample preparation & xenograft surgery procedure

LAPC9 and BM18 xenografts were maintained subcutaneously in 6 week old, CB17 SCID male mice, under anesthesia (Domitor® 0.5mg/kg, Dormicum 5mg/kg, Fentanyl 0.05mg/kg). All animal experiments were approved by ethical committee of Canton Bern (animal license BE55/16, BE12/17). Castration was achieved by bilateral orchiectomy. For androgen replacement, testosterone propionate dissolved in castor oil (Sigma, 86541-5G) was administered by single subcutaneous injection (2mg per dosage, 25G needle).

### RNA isolation from tissue samples

Tissue RNA was extracted using standard protocol of Qiazol (Qiagen) tissue lysis by TissueLyser (2min, 20Hz). Quality of RNA was assessed by Bioanalyzer (Agilent). RNA from FFPE material was extracted using the Maxwell® 16 LEV RNA FFPE Purification Kit (Promega, AS1260).

### RNA sequencing

RNA extracted from BM18, LAPC9 whole PDX tumor extracts (300ng) were subjected to RNA sequencing. Specimens were prepared for RNA sequencing using TruSeq RNA Library Preparation Kit v2 or riboZero as previously described [41]. RNA integrity was verified using the Agilent Bioanalyzer 2100 (Agilent Technologies). cDNA was synthesized from total RNA using Superscript III (Invitrogen). Sequencing was then performed on GAII, HiSeq 2000, or HiSeq 2500. For the RNASeq, the NEBNext Ultra II Directional RNA Library Prep Kit for Illumina was used to process the sample(s). The sample preparation was performed according to the protocol “NEBNext Ultra II Directional RNA Library Prep Kit for Illumina” (NEB #E7760S/L). Briefly, mRNA was isolated from total RNA using the oligo-dT magnetic beads. After fragmentation of the mRNA, a cDNA synthesis was performed. This was used for ligation with the sequencing adapters and PCR amplification of the resulting product. The quality and yield after sample preparation was measured with the Fragment Analyzer. The size of the resulting products was consistent with the expected size distribution (a broad peak between 300-500 bp). Clustering and DNA sequencing using the NovaSeq6000 was performed according to manufacturer’s protocols. A concentration of 1.1 nM of DNA was used. Image analysis, base calling, and quality check was performed with the Illumina data analysis pipeline RTA3.4.4 and Bcl2fastq v2.20. Sequence reads were aligned using STAR two-pass to the human reference genome GRCh37 [42], and mouse reference genome GRCm38. Gene counts were quantified using the “GeneCounts” option. Per-gene counts-per-million (CPM) were computed and log_2_-transformed adding a pseudo-count of 1 to avoid transforming 0. Genes with log_2_-CPM <1 in more than three samples were removed. Differential expression analysis was performed using the edgeR package [43]. Normalization was performed using the “TMM” (weighted trimmed mean) method and differential expression was assessed using the quasi-likelihood F-test. Genes with FDR <0.05 and > 2-fold were considered significantly differentially expressed. RSEM was used to obtain TPM (Transcript per million) counts.

### Tissue dissociation and MACS

Tumor tissue is collected in Basis medium (Advanced DMEM F12 Serum Free medium (Thermo, 12634010) containing 10mM Hepes (Thermo,15630080), 2mM GlutaMAX supplement (Thermo, 35050061) and 100 μg/ml Primocin (InVivoGen, ant-pm-1)). After mechanical disruption the tissue is washed in Basis medium (220rcf, 5min) and incubated in enzyme mix for tissue dissociation (collagenase type II enzyme mix (Gibco, 17101-015, 5mg/ml dissolved in Basis medium, DNase: 15ug/ml (Roche, 10104159001) and 10 μM Y-27632-HCl Rock inhibitor (Selleckchem, S1049). Enzyme mix volume is adjusted so that the tissue volume does not exceed 1/10 of the total volume and tissue is incubated at 37°C for 1-2h with mixing every 20 minutes. After digestion of large pieces is complete, the suspension is passed through 100um cell strainer (Falcon®, VWR 734-0004) attached to a 50ml Falcon tube. Using a syringe rubber to crash tissue against the strainer & wash in 5ml basic medium (220rcf, 5min). Cell pellet is incubated in 5ml precooled EC lysis buffer (*150 mM NH_4_Cl, 10 mM KHCO_3_, 0.1mM EDTA)*, incubated for 10 min, washed in equal volume of basis medium followed by centrifugation (220rcf, 5min). Pellet is resuspended in 2-5ml accutase™ (StemCell Technologies, 07920), depending on the sample amount; biopsies vs tissue and incubated for 10min at room temperature. The cell suspension is passed through 40um pore size strainer (Falcon®, VWR 734-0004), and the strainer is washed by adding 2ml of accutase on the strainer. Single cell suspension is counted to determine seeding density, and washed in 5ml of basis medium and spin down 220rcf, 5min. Magnetic cell sorting was performed to separate purified human versus mouse cell fractions using the Mouse Cell Depletion Kit (Miltenyi Biotek, 130-104-694). For the proteomic experiments, cell fractions from tumor tissues (N=3-4 per condition) were pooled together in order to suffice for 1×10^6 cells, representing one technical replicate per sample.

### Proteomics

#### Sample preparation

Approx. 1×10^6 cell pellets (N=1 technical replicate per condition deriving from N=3-4 biological replicate samples) were resuspended in 50 μl PBS following the addition of 50 μl 1% SDS in 100 mM Hepes/NaOH pH8.5 supplemented with protease inhibitor cocktail EDTA-free (Roche). Samples were heated to 95°C for 5 min, transferred on ice and benzonase (Merck, #71206-3) was added to degrade DNA at 37°C for 30 minutes. Samples were reduced by the addition of 2 μl of a 200 mM DTT solution in 200 mM Hepes/NaOH pH 8.5 and subsequently alkylated by the addition of 4 μl of a 400 mM chloroacetamide (CAA, Sigma-Aldrich, #C0267) solution in 200 mM Hepes/NaOH pH 8.5. Samples were incubated at 56°C for 30 min. Access CAA was quenched by the addition of 4 μl of a 200 mM DTT solution in 200 mM Hepes/NaOH pH 8.5. Lysate were subjected to an in-solution tryptic digest using the Single-Pot Solid-Phase-enhanced Sample Preparation (SP3) protocol [44, 45]. To this end, 20 μl of Sera-Mag Beads (Thermo Scientific, #4515-2105-050250, 6515-2105-050250) were mixed, washed with H_2_O and resuspended in 100 μl H_2_O. 2 μl of freshly prepared bead mix and 5 μl of an aqueous 10% formic acid were added to 40 μl of lysates to achieve an acidic pH. 47 μl acetonitrile were added and samples were incubated for 8 min at room temperature. Beads were captured on a magnetic rack and washed three times with 70% ethanol and once with acetonitrile. 0.8 μg of sequencing grade modified trypsin (Promega, #V5111) in 10 μl 50 mM Hepes/NaOH pH8.5 were added. Samples were digested over night at 37°C. Beads were captured and the supernatant transferred and dried down. Peptides were reconstituted in 10 μl of H_2_O and reacted with 80 μg of TMT10plex (Thermo Scientific, #90111) [46] label reagent dissolved in 4 μl of acetonitrile for 1 h at room temperature. Excess TMT reagent was quenched by the addition of 4 μl of an aqueous solution of 5% hydroxylamine (Sigma, 438227). Mixed peptides were subjected to a reverse phase clean-up step (OASIS HLB 96-well μElution Plate, Waters #186001828BA) and analyzed by LC-MS/MS on a Q Exactive Plus (Thermo Scentific) as previously described [47].

#### Mass spectrometric analysis

Briefly, peptides were separated using an UltiMate 3000 RSLC (Thermo Scientific) equipped with a trapping cartridge (Precolumn; C18 PepMap 100, 5 lm, 300 lm i.d. × 5 mm, 100 A°) and an analytical column (Waters nanoEase HSS C18 T3, 75 lm × 25 cm, 1.8 lm, 100 A°). Solvent A: aqueous 0.1% formic acid; Solvent B: 0.1% formic acid in acetonitrile (all solvents were of LC-MS grade). Peptides were loaded on the trapping cartridge using solvent A for 3 min with a flow of 30 μl/min. Peptides were separated on the analytical column with a constant flow of 0.3 μl/min applying a 2 h gradient of 2 – 28% of solvent B in A, followed by an increase to 40% B. Peptides were directly analyzed in positive ion mode applying with a spray voltage of 2.3 kV and a capillary temperature of 320°C using a Nanospray-Flex ion source and a Pico-Tip Emitter 360 lm OD × 20 lm ID; 10 lm tip (New Objective). MS spectra with a mass range of 375–1.200 m/z were acquired in profile mode using a resolution of 70.000 [maximum fill time of 250 ms or a maximum of 3e6 ions (automatic gain control, AGC)]. Fragmentation was triggered for the top 10 peaks with charge 2–4 on the MS scan (data-dependent acquisition) with a 30 second dynamic exclusion window (normalized collision energy was 32). Precursors were isolated with a 0.7 m/z window and MS/MS spectra were acquired in profile mode with a resolution of 35,000 (maximum fill time of 120 ms or an AGC target of 2e5 ions).

#### Raw MS Data Analysis

Acquired data were analyzed using IsobarQuant [48] and Mascot V2.4 (Matrix Science) using either a reverse UniProt FASTA Mus musculus (UP000000589) or Homo sapiens (UP000005640) database. Moreover, a combined database thereof has been generated and used for the analysis. These databases also included common contaminants. The following modifications were taken into account: Carbamidomethyl (C, fixed), TMT10plex (K, fixed), Acetyl (N-term, variable), Oxidation (M, variable) and TMT10plex (N-term, variable). The mass error tolerance for full scan MS spectra was set to 10 ppm and for MS/MS spectra to 0.02 Da. A maximum of 2 missed cleavages were allowed. A minimum of 2 unique peptides with a peptide length of at least seven amino acids and a false discovery rate below 0.01 were required on the peptide and protein level [49].

#### MS Data Analysis

The raw output files of IsobarQuant (protein.txt – files) were processed using the R programming language (ISBN 3-900051-07-0). As a quality filter, only proteins were allowed that were quantified with at least two unique peptides. Human and mouse samples were searched against a combined human and mouse data base and annotated as unique for human or mouse or mixed. Raw signal-sums (signal_sum columns) were normalized using vsn (variance stabilization normalization [50]. In order to try to annotate each observed ratio with a p-value, each ratio distribution was analyzed with the locfdr function of the locfdr package [51] to extract the average and the standard deviation (using maximum likelihood estimation). Then, the ratio distribution was transformed into a z-distribution by normalizing it by its standard deviation and mean. This z-distribution was analyzed with the fdrtool function of the fdrtool package [52] in order to extract p-values and false discovery rates (fdr - q-values).

#### Tissue microarray

Kaplan Meier curves to calculate the association between TNC-positive cells and disease progression were calculated using the “survfit” function and the global Log-Rank test using the Survival R package [53, 54]. To estimate survival we used the function “surv_cutpoint” which employs maximally selected rank statistics (maxstat) to determine the optimal cutpoint for continous variables [18]. For pairwise comparison, p-value was estimated by the Log-Rank test and adjusted with Benjamini–Hochberg (BH) method. If no information on patient outcome was available, information at last follow-up was used for all parameters. Clinical progression was defined as metastasis or local recurrence. Disease progression was defined by combining any form of recurrence (PSA and clinical progression). Data representation and graphical plots were generated using the ggplot2 R package [55]. Data analyses were done using RStudio version 1.1.463 [56] and R version 3.5.3 [57].

#### Fibroblast cultures

LAPC9 fibroblasts were derived by tumor tissue outgrowth in DMEM high glucose (Thermo, 61965059) supplemented with 10% fetal calf serum, 1% penicillin-streptomycin (Sigma), 1% Insulin Transferin Selenium (Thermo, 41400045) and 10ng/ml FGF2 (Peprotech, 100-18B). Following cell outgrowth and expansion, mouse stromal cells were purified by MACS depletion and maintained in the media described.

#### Immunohistochemistry

FFPE sections (4um) were deparaffinised and used for heat mediated antigen retrieval (citrate buffer pH 6, Vector labs). Sections were blocked for 10min in 3%H2O2, followed by 30min, RT incubation in 1%BSA in PBS-0.1%Tween20. The following antibodies were used:

| Dilution | Antibody | Company | Catalog No |
| --- | --- | --- | --- |
| 1to500 | **Ki67** | Gene Tex | GTX16667 |
| 1to100 | **Tnc** | abcam | ab133273 |

Secondary anti-rabbit antibody Envision HRP (DAKO) for 30min or anti-rat HRP (Thermo). Signal detection with AEC substrate (DAKO). Sections were counterstained with Hematoxylin and mounted with Aquatex.

## ACKNOWLEDGEMENTS

This project has received funding from the European Union’s Horizon 2020 research and innovation programme under the Marie Skłodowska-Curie grant agreement No 748836 (STOPCa).

## REFERENCES

1. Heidenreich, A., et al., EAU guidelines on prostate cancer. Part II: Treatment of advanced, relapsing, and castration-resistant prostate cancer. Eur Urol, 2014. 65(2): p. 467–79.

2. Malanchi, I., et al., Interactions between cancer stem cells and their niche govern metastatic colonization. Nature, 2011. 481(7379): p. 85–9.

3. Shiozawa, Y., et al., Human prostate cancer metastases target the hematopoietic stem cell niche to establish footholds in mouse bone marrow. The Journal of Clinical Investigation, 2011. 121(4): p. 1298–1312.

4. Hensel, J., et al., Osteolytic cancer cells induce vascular/axon guidance processes in the bone/bone marrow stroma. Oncotarget, 2018. 9(48).

5. Ozdemir, B.C., et al., The molecular signature of the stroma response in prostate cancer-induced osteoblastic bone metastasis highlights expansion of hematopoietic and prostate epithelial stem cell niches. PLoS One, 2014. 9(12): p. e114530.

6. Rucci, N. and A. Teti, Osteomimicry: How the Seed Grows in the Soil. Calcified Tissue International, 2018. 102(2): p. 131–140.

7. Tyekucheva, S., et al., Stromal and epithelial transcriptional map of initiation progression and metastatic potential of human prostate cancer. Nature Communications, 2017. 8(1): p. 420.

8. Setlur, S.R. and M.A. Rubin, Current thoughts on the role of the androgen receptor and prostate cancer progression. Adv Anat Pathol, 2005. 12(5): p. 265–70.

9. Leach, D.A., et al., Stromal androgen receptor regulates the composition of the microenvironment to influence prostate cancer outcome. Oncotarget, 2015. 6(18).

10. Leach, D.A., et al., Cell-lineage specificity and role of AP-1 in the prostate fibroblast androgen receptor cistrome. Molecular and Cellular Endocrinology, 2017. 439: p. 261–272.

11. Nash, C., et al., Genome-wide analysis of AR binding and comparison with transcript expression in primary human fetal prostate fibroblasts and cancer associated fibroblasts. Mol Cell Endocrinol, 2018. 471: p. 1–14.

12. Thalmann, G.N., et al., Human prostate fibroblasts induce growth and confer castration resistance and metastatic potential in LNCaP Cells. European urology, 2010. 58(1): p. 162–171.

13. Thalmann, G.N., et al., Androgen-independent cancer progression and bone metastasis in the LNCaP model of human prostate cancer. Cancer Res, 1994. 54(10): p. 2577–81.

14. Briganti, A., et al., Impact of age and comorbidities on long-term survival of patients with high-risk prostate cancer treated with radical prostatectomy: a multi-institutional competing-risks analysis. Eur Urol, 2013. 63(4): p. 693–701.

15. Tosco, L., et al., The EMPaCT Classifier: A Validated Tool to Predict Postoperative Prostate Cancer-related Death Using Competing-risk Analysis. Eur Urol Focus, 2018. 4(3): p. 369–375.

16. Chys, B., et al., Preoperative Risk-Stratification of High-Risk Prostate Cancer: A Multicenter Analysis. Frontiers in Oncology, 2020. 10(246).

17. D’Amico, A.V., et al., Biochemical Outcome After Radical Prostatectomy, External Beam Radiation Therapy, or Interstitial Radiation Therapy for Clinically Localized Prostate Cancer. JAMA, 1998. 280(11): p. 969–974.

18. Kassambara, A., Survminer: Drawing Survival Curves using ‘ggplot2’. In., 0.4.3 edn. 2018.

19. Sahai, E., et al., A framework for advancing our understanding of cancer-associated fibroblasts. Nature reviews. Cancer, 2020. 20(3): p. 174–186.

20. Corn, P.G., The tumor microenvironment in prostate cancer: elucidating molecular pathways for therapy development. Cancer management and research, 2012. 4: p. 183–193.

21. Bissell, M.J. and W.C. Hines, Why don’t we get more cancer? A proposed role of the microenvironment in restraining cancer progression. Nature medicine, 2011. 17(3): p. 320–329.

22. Petersen, O.W., et al., Interaction with basement membrane serves to rapidly distinguish growth and differentiation pattern of normal and malignant human breast epithelial cells. Proceedings of the National Academy of Sciences of the United States of America, 1992. 89(19): p. 9064–9068.

23. Weaver, V.M., et al., Reversion of the malignant phenotype of human breast cells in three-dimensional culture and in vivo by integrin blocking antibodies. The Journal of cell biology, 1997. 137(1): p. 231–245.

24. Sung, S.-Y., et al., Coevolution of Prostate Cancer and Bone Stroma in Three-Dimensional Coculture: Implications for Cancer Growth and Metastasis. Cancer Research, 2008. 68(23): p. 9996–10003.

25. Cutz, J.-C., et al., Establishment in Severe Combined Immunodeficiency Mice of Subrenal Capsule Xenografts and Transplantable Tumor Lines from a Variety of Primary Human Lung Cancers: Potential Models for Studying Tumor Progression–Related Changes. Clinical Cancer Research, 2006. 12(13): p. 4043–4054.

26. Hidalgo, M., et al., Patient-derived xenograft models: an emerging platform for translational cancer research. Cancer discovery, 2014. 4(9): p. 998–1013.

27. Nash, C., et al., Genome-wide analysis of AR binding and comparison with transcript expression in primary human fetal prostate fibroblasts and cancer associated fibroblasts. Molecular and Cellular Endocrinology, 2018. 471: p. 1–14.

28. Tuxhorn, J.A., et al., Reactive Stroma in Human Prostate Cancer. Induction of Myofibroblast Phenotype and Extracellular Matrix Remodeling, 2002. 8(9): p. 2912–2923.

29. Ibrahim, S.N., et al., Tenascin expression in prostatic hyperplasia, intraepithelial neoplasia, and carcinoma. Human Pathology, 1993. 24(9): p. 982–989.

30. Xue, Y., et al., Tenascin-C expression in prostatic intraepithelial neoplasia (PIN): a marker of progression? Anticancer Res, 1998. 18(4A): p. 2679–84.

31. Ni, W.-D., et al., Tenascin-C is a potential cancer-associated fibroblasts marker and predicts poor prognosis in prostate cancer. Biochemical and Biophysical Research Communications, 2017. 486(3): p. 607–612.

32. Cai, M., et al., Degradation of Tenascin-C and Activity of Matrix Metalloproteinase-2 Are Associated with Tumor Recurrence in Early Stage Non-Small Cell Lung Cancer. Clinical Cancer Research, 2002. 8(4): p. 1152–1156.

33. Kusagawa, H., et al., Expression and degeneration of tenascin-C in human lung cancers. British Journal of Cancer, 1998. 77(1): p. 98–102.

34. San Martin, R., et al., Tenascin-C and Integrin α9 Mediate Interactions of Prostate Cancer with the Bone Microenvironment. Cancer Research, 2017. 77(21): p. 5977–5988.

35. Colombel, M., et al., Increased expression of putative cancer stem cell markers in primary prostate cancer is associated with progression of bone metastases. Prostate, 2012. 72(7): p. 713–20.

36. Lowy, C.M. and T. Oskarsson, Tenascin C in metastasis: A view from the invasive front. Cell Adhesion & Migration, 2015. 9(1-2): p. 112–124.

37. Mishra, P., et al., Genomic alterations of Tenascin C in highly aggressive prostate cancer: a meta-analysis. Genes & cancer, 2019. 10(5-6): p. 150–159.

38. Kiebish, M.A., et al., Multi-omic serum biomarkers for prognosis of disease progression in prostate cancer. Journal of translational medicine, 2020. 18(1): p. 10–10.

39. Mo, F., et al., Stromal Gene Expression is Predictive for Metastatic Primary Prostate Cancer. European urology, 2018. 73(4): p. 524–532.

40. Jachetti, E., et al., Tenascin-C Protects Cancer Stem–like Cells from Immune Surveillance by Arresting T-cell Activation. Cancer Research, 2015. 75(10): p. 2095–2108.

41. Beltran, H., et al., Whole-Exome Sequencing of Metastatic Cancer and Biomarkers of Treatment Response. JAMA Oncol, 2015. 1(4): p. 466–74.

42. Dobin, A., et al., STAR: ultrafast universal RNA-seq aligner. Bioinformatics (Oxford, England), 2013. 29(1): p. 15–21.

43. Nikolayeva, O. and M.D. Robinson, edgeR for differential RNA-seq and ChIP-seq analysis: an application to stem cell biology. Methods Mol Biol, 2014. 1150: p. 45–79.

44. Hughes, C.S., et al., Ultrasensitive proteome analysis using paramagnetic bead technology. Mol Syst Biol, 2014. 10: p. 757.

45. Moggridge, S., et al., Extending the Compatibility of the SP3 Paramagnetic Bead Processing Approach for Proteomics. J Proteome Res, 2018. 17(4): p. 1730–1740.

46. Werner, T., et al., Ion coalescence of neutron encoded TMT 10-plex reporter ions. Anal Chem, 2014. 86(7): p. 3594–601.

47. Becher, I., et al., Pervasive Protein Thermal Stability Variation during the Cell Cycle. Cell, 2018. 173(6): p. 1495–1507 e18.

48. Franken, H., et al., Thermal proteome profiling for unbiased identification of direct and indirect drug targets using multiplexed quantitative mass spectrometry. Nat Protoc, 2015. 10(10): p. 1567–93.

49. Savitski, M.M., et al., A Scalable Approach for Protein False Discovery Rate Estimation in Large Proteomic Data Sets. Mol Cell Proteomics, 2015. 14(9): p. 2394–404.

50. Huber, W., et al., Variance stabilization applied to microarray data calibration and to the quantification of differential expression. Bioinformatics, 2002. 18 Suppl 1: p. S96–104.

51. Efron, B., Large-Scale Simultaneous Hypothesis Testing. Journal of the American Statistical Association, 2004. 99(465): p. 96–104.

52. Strimmer, K., fdrtool: a versatile R package for estimating local and tail area-based false discovery rates. Bioinformatics, 2008. 24(12): p. 1461–2.

53. Therneau, T., A Package for Survival Analysis in S. In., 2.38 edn;. 2015.

54. Therneau, T., PMG: modeling survival data: extending the cox model. In. New York: Springer;. 2000.

55. Wickham, H., ggplot2: elegant graphics for data analysis. In. New York: Springer-Verlag. 2016.

56. RStudio Team: RStudio: integrated development for R. RStudio. In. Boston, M.I., 2016.

57. R Core Team: R: a language and environment for statistical computing. In. Vienna, A.R.F.f.S.C., 2019.

